# Musashi proteins are post-transcriptional regulators of the epithelial-luminal cell state

**DOI:** 10.1101/006270

**Authors:** Yarden Katz, Feifei Li, Nicole Lambert, Ethan Sokol, Wai-Leong Tam, Albert W. Cheng, Edoardo M. Airoldi, Christopher J. Lengner, Piyush B. Gupta, Zhengquan Yu, Rudolf Jaenisch, Christopher B. Burge

## Abstract

**Summary:** The conserved Musashi (Msi) family of RNA binding proteins are expressed in stem/progenitor and cancer cells, but mostly absent from differentiated cells, consistent with a role in cell state regulation. We found that Msi genes are rarely mutated but frequently overexpressed in human cancers, and associated with an epithelial-luminal cell state. Using ribosome footprint profiling and RNA-seq analysis of genetic mouse models in neuronal and mammary cell types, we found that Msis regulate translation of genes implicated in epithelial cell biology and epithelial-to-mesenchymal transition (EMT) and promote an epithelial splicing pattern. Overexpression of Msi proteins inhibited translation of genes required for EMT, including Jagged1, and repressed EMT in cell culture and in mammary gland *in vivo*, while knockdown in epithelial cancer cells led to loss of epithelial identity. Our results show that mammalian Msi proteins contribute to an epithelial gene expression program and promote an epithelial-luminal state in both neural and breast cell types.

**Highlights:**

- Msi proteins bind UAG motifs in vitro and in 3’ UTRs of mRNAs
- Msi proteins are markers of epithelial state in brain and breast tumors, and cell lines
- The Notch regulator *Jag1* mRNA is bound and translationally repressed by Msi
- Msi overexpression represses EMT in the mammary gland *in vivo*

## Introduction

During both normal development and cancer progression, cells undergo state transitions marked by distinct gene expression profiles and changes in morphology, motility and other properties. The Epithelial-to-Mesenchymal Transition (EMT) is one such transition, which is essential in development and is thought to be recruited by tumor cells undergoing metastasis (Polyak and Weinberg, 2009). Much work on cell state transitions in both the stem cell and cancer biology fields has focused on the roles that transcription factors play in driving these transitions (Lee and Young, 2013; Polyak and Weinberg, 2009), such as the induction of EMT by ectopic expression of the transcription factors Snail, Slug or Twist (Mani et al., 2008).

Recent work has shown that RNA-binding proteins (RBPs) also play important roles in cell state transitions, by driving post-transcriptional gene expression programs specific to a particular cell state. The epithelial specific regulatory protein (ESRP) family of RBPs are RNA splicing factors with epithelial tissue-specific expression whose ectopic expression can partially reverse EMT (Shapiro et al., 2011; Warzecha et al., 2009). RBPs have also been implicated in other cell state transitions, such as reprogramming of somatic cells to induced pluripotent stem cells (iPSCs), which have the essential characteristics of embryonic stem cells (ESCs). For example, overexpression of the translational regulator and miRNA processing factor Lin28 along with three transcription factors is sufficient to reprogram somatic cells (Yu et al., 2007). The Muscleblind-like (Mbnl) family of RBPs promote differentiation by repressing an ESC-specific alternative splicing program, and inhibition of Mbnls promotes reprogramming (Han et al., 2013). For ESRP, Lin28 and Mbnl proteins, the developmental or cell-type-specific expression pattern of the protein provided clues to their functions in maintenance of epithelial, stem cell or differentiated cell state.

The Musashi (Msi) family comprises some of the most highly conserved and tissue-specific RBPs, with Msi in *Drosophila* expressed exclusively in the nervous system (Busch and Hertel, 2011; Nakamura et al., 1994). In mammals, the two family members Msi1 and Msi2 are highly expressed in stem cell compartments but are mostly absent from differentiated tissues. Msi1 is a marker of neural stem cells (NSCs) (Sakakibara et al., 1996) and is also expressed in stem cells in the gut (Kayahara et al., 2003) and epithelial cells in the mammary gland (Colitti and Farinacci, 2009), while Msi2 is expressed in hematopoietic stem cells (HSCs) (Kharas et al., 2010). This expression pattern led to the proposal that Msi proteins generally mark the epithelial stem cell state across distinct tissues (Okano et al., 2005), with HSCs being an exception. Msi1 is not expressed in the normal adult brain outside a minority of adult NSCs, but is induced in glioblastoma (Muto et al., 2012).

Msi proteins affect cell proliferation in several cancer types. In glioma and medulloblastoma cell lines, knockdown of Msi1 reduced the colony forming capacity of these cells and reduced their tumorigenic growth in a xenograft assays in mice (Muto et al., 2012). Msi expression correlates with HER2 expression in breast cancer cell lines, and knockdown of Msi proteins resulted in decreased proliferation (Wang et al., 2010). These data, coupled with the cell-type specific expression of Msi proteins in normal development, suggested that Msi proteins might function as regulators of cell state, with potential relevance to cancer cell state.

Msi proteins have been proposed to act as translational repressors of mRNAs (and sometimes as activators (MacNicol et al., 2011)) through binding of mRNA 3’ UTRs, and were speculated to affect pre-mRNA processing in *Drosophila* (Nakamura et al., 1994; Okano et al., 2002). However, no conclusive genome-wide evidence for either role for the mammalian Msi family has been reported. We aimed to investigate the roles of these proteins in human cancers, and gain a better understanding of their genome-wide effects on the transcriptome.

## Results

### Msi genes are frequently overexpressed in multiple human cancers

To obtain a broad view of the role Msis might play in human cancer, we surveyed the expression and mutation profiles of Msi genes in primary tumors using genome and RNA sequencing data (RNA-Seq) from The Cancer Genome Atlas (TCGA) (TCGA, 2012). To determine whether Msi genes are generally upregulated in human cancers, we analyzed RNA-Seq data from 5 cancer types for which matched tumor-control pairs were available. In these matched designs, a pair of RNA samples are obtained in parallel from a single patient’s tumor and healthy tissue-matched biopsy, thus minimizing the contribution of individual genetic variation to expression differences. We observed that Msi1 was upregulated in at least 40% of breast, lung and prostate tumors, while Msi2 was upregulated in at least 50% of breast and prostate tumors (**Figure 1A**, top). Overall, Msi1 or Msi2 were significantly upregulated in matched tumor-control pairs of 3 of the 5 cancer types, compared to control pairs. Kidney tumors showed the opposite expression pattern, with Msi1 and Msi2 downregulated in a majority of tumors and rarely upregulated, and in thyroid cancer neither Msi1 nor Msi2 showed a strong bias towards up- or down-regulation (**Figure 1A**, top). In breast tumors, a bimodal distribution of Msi1 expression was observed, with a roughly even split between up- and down-regulation of Msi1, consistent with the idea that Msi1 upregulation might be specific to a subtype of breast tumors. The bimodality of Msi1 expression was not seen when comparing control pairs, so is not explained by general variability in Msi1 levels (**Figure 1A**, bottom, solid versus dotted lines).

**Figure 1.**
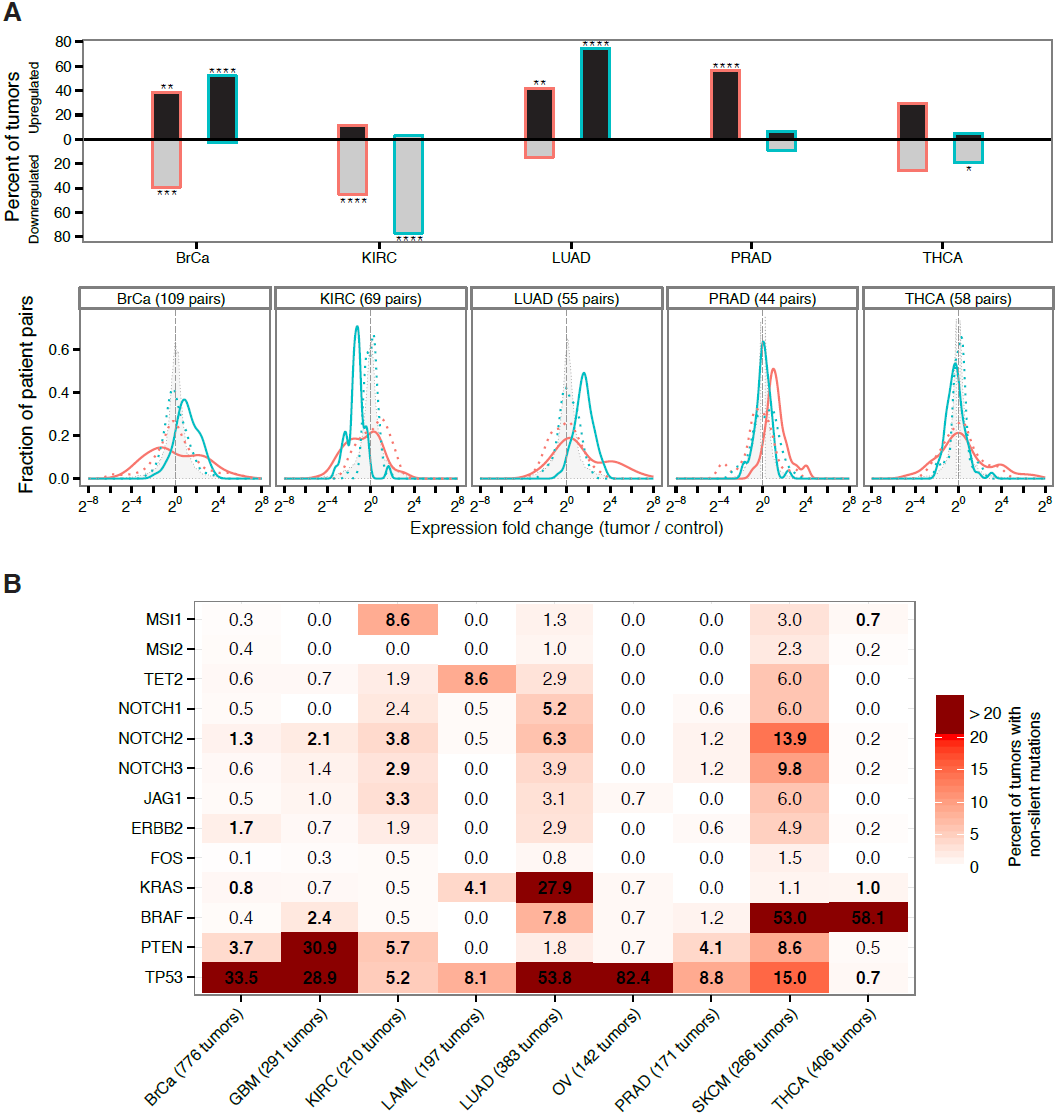
Msi genes are frequently overexpressed in breast, lung and prostate cancer but downregulated in kidney cancer. **(A)** Top: Percentage of matched tumor-control pairs with upregulated (grey-fill bars) or downregulated (black-fill bars) Msi1 or Msi2 in 5 cancer types with matched RNA-Seq data. Upregulated/downregulated defined as ≥ 2-fold change in expression in tumor relative to matched control. Asterisks indicate one-tailed statistical significance levels relative to control pairs. Bottom: Distribution of fold changes for Msi1 and Msi2 in matched tumor-control pairs (solid red and green lines, respectively) and in an equal number of control pairs (dotted red and green lines, respectively.) Shaded gray density shows the fold change across all genes. **(B)** Percentage of tumors with non-silent mutations in Msi1/Msi2 and a select set of oncogenes and tumor suppressors across 9 cancer types. Bold entries indicate genes whose mutation rate is at least two-fold above the cancer type average mutation rate.

Examining genome sequencing data from matched tumor-control pairs across 9 diverse cancer types, we found that Msi1 and Msi2 were not significantly mutated in majority of these cancers (**Figure 1B**). One notable exception was kidney cancer (KIRC) where non-silent mutations in Msi1 were present in nearly 9% of tumors, far exceeding the background mutation rate of genes in kidney tumors (**Supp. Figure 1A**). This observation, together with the lower Msi mRNA levels observed in matched kidney tumors (**Figure 1A**), is consistent with a model in which loss of Msi function is selected for in kidney tumor cells, either as a result of downregulation or non-silent mutations. The observation Msi1/Msi2 were not significantly mutated in most tumors but are overexpressed in several tumor typess (including glioblastoma) makes their profile more similar to oncogenes like FOS or HER2, than to tumor suppressors like PTEN and TP53, which tend to have the opposite pattern (**Figure 1B**).

### Msi expression marks an epithelial-luminal state and is downregulated upon EMT

To determine whether Msi overexpression is specific to a particular cancer cell state, we focused on breast cancer, where tumors with distinct properties can be robustly classified by gene expression (Parker et al., 2009; TCGA, 2012). Unsupervised hierarchical clustering of matched tumor and control samples produced a nearly perfect separation of tumors from control samples, rather than clustering by patient/genome of origin (**Supp. Figure 1B**). We overlaid on top of our clustering a classification of samples into Normal, HER2+, Luminal A, Luminal B and Basal states using RNA-Seq expression of the PAM50 gene set (Parker et al., 2009). Our clustering using all genes corresponded well to the PAM50 classification (TCGA, 2012), separating most Luminal A from Luminal B tumors and showing a general grouping of HER2+ tumors (**Supp. Figure 1B**). Using this classification, we found that Msi2 was highly expressed in Luminal tumors (**Figure 2A**). Msi1 was more variable across tumor subtypes, often showing a bimodal profile split between up- and down-regulation (**Figure 1A** and **Supp. Figure 1B**). Msi2 expression was highest in Luminal B tumors, which are known to be more aggressive and highly proliferating (Ki67-high) than Luminal A types, and are thought to share properties with epithelial mammary progenitors cells (Das et al., 2013). These observations prompted the hypothesis that Msi proteins might be localized to epithelial cells in breast cancer tumors.

**Figure 2.**
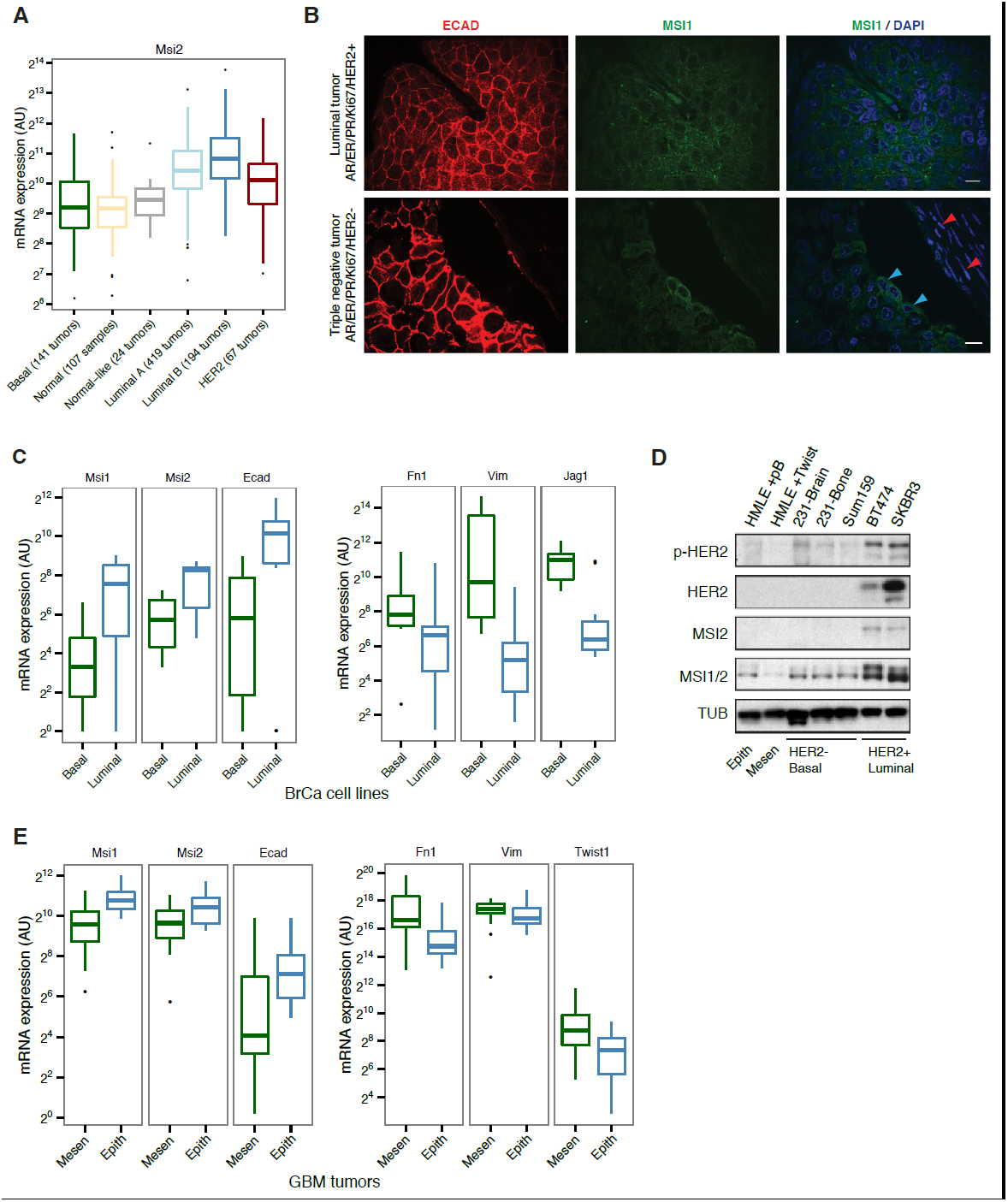
Msi is associated with the epithelial-luminal state in breast cancer. **(A)** mRNA expression of Msi2 by RNA-Seq across different breast tumor types in TCGA RNA-Seq. **(B)** Immunofluorescence staining for Ecadherin (red) and Msi1 (green). Top: luminal human breast tumor with high number of ECAD-positive cells. MSI1 shows primarily cytoplasmic localization (white arrowheads). Bottom: Triple negative, basal-like tumor. ECAD-positive cells showed strong cytoplasmic MSI1 stain (blue arrowheads) while ECAD-negative cells were MSI1-negative (red). Single confocal stacks shown, 10 μm scale. **(C)** mRNA expression of Msi1, Msi2, Ecad, Fn1, Vim and Jag1 in breast cancer cell lines by RNA-Seq (**Supp. Table 1** describes datasets). **(D)** Western blot for MSI1/2 (MSI1/2 cross react. antibody), MSI2, phosphorylated HER2 (p-HER2) and HER2 in panel of breast cell lines. ‘HMLE + pB’ indicates HMLE cells infected with pB empty vector, ‘HMLE + Twist’ indicates HMLE cells infected with Twist transcription factor to induce EMT. MDAMB231-derived metastatic lines (231-Brain, 231-Bone) and Sum159 are basal, HER2-negative cancer cell lines. BT474 and SKBR3 are HER2-positive, epithelial-luminal cancer cell lines. Epithelial-luminal (HER2-positive) lines show increased expression of Msi proteins compared with basal lines, and Twist-induced EMT reduces Msi expression. **(E)** mRNA expression of Msi1, Msi2, Ecad, Fn1, Vim and Twist1 in GBM tumors classified as mesenchymal (*n* = 20) or epithelial (*n* = 20) using an EMT gene signature.

To examine expression and distribution of Msi proteins in tumors, we stained a panel of human breast cancer tumors for MSI1 and the epithelial marker E-cadherin (ECAD). MSI1 expression was predominantly cytoplasmic (**Figure 2B**, top panel). Across luminal tumors, MSI1 was co-expressed with ECAD (as in **Figure 2B**, top panel). In triple negative/basal-like tumors, the minority of ECAD-positive cells showed strong MSI1 staining, whereas ECAD-negative cells showed little to no expression (**Figure 2B**, blue and red arrowheads, respectively), consistent with the observation that Msi is associated with an epithelial cell state in tumors.

To explore whether Msi expression is associated with a luminal as opposed to basal state in a more homogenous system, we collected RNA-Seq data for luminal and basal breast cancer cell lines generated by multiple independent labs (see **Supp. Table 1** for RNA-Seq datasets used). Gene expression profiles from the same cell lines generated independently tended to cluster together in unsupervised clustering (supporting the use of cross-lab comparisons), and overall the basal cell lines were distinguishable from the luminal lines (**Supp. Figure 2A**). Matching the pattern observed in primary tumors, we observed higher Msi1 and Msi2 expression in luminal breast cancer lines than in basal lines (**Figure 2C**, left panel). Fibronectin (Fn1), Vimentin (Vim) and Jagged 1 (Jag1) which are associated with the basal/mesenchymal state (Yamamoto et al., 2013) had the opposite pattern, showing ∼60-fold enrichment in basal over luminal lines (**Figure 2C**, right panel). The enrichments of these four genes for either the luminal or basal state were unusual when compared to the background distribution of these enrichments across all expressed genes (**Supp. Figure 2C**), indicating that these genes are strong indicators of the two states.

To further investigate the connection between Msi expression and EMT in breast cancer, we examined Msi expression in a panel of breast cancer-derived cell lines. Consistent with the RNA-Seq data from primary tumors, HER2+ epithelial cell lines expressed higher levels of Msi1 and Msi2 compared with HER2– lines (**Figure 2D**, lane 6 and 7). A standard cell culture model of EMT is the immortalized inducible-Twist human mammary epithelial (HMLE-Twist) cell line, which undergoes EMT when induced to express the transcription factor Twist (Mani et al., 2008). We found that Msi1 was strongly downregulated in HMLE cells following Twist-induced EMT (**Figure 2D**), consistent with the epithelial-associated expression pattern of Msi in primary tumors (**Figure 2A-C**). Similarly, Msi protein expression was higher in luminal, HER2+ breast cancer lines (BT474, SKBR3 in **Figure 2D**) compared with basal HER2– breast cancer lines (brain and bone metastatic derivatives of MDAMB231, 231-Brain and 231-Bone, and SUM159 in **Figure 2D**).

We next wanted to test whether the epithelial expression signature of Msis is present in other primary tumors. Given the established role of Msi proteins as regulators of Glioblastoma (GBM) cell growth and as markers of primary tumors (Muto et al., 2012), we next asked whether there is a similar subtype expression pattern in GBM tumors from TCGA (Verhaak et al., 2010). We used an EMT gene signature to rank GBM tumors as more epithelial or more mesenchymal, based on the similarity of each tumor’s gene expression profile to that of cells undergoing EMT in culture (Feng et al., 2014). Using this ranking, we found that the top 20 most mesenchymal tumors expressed lower levels of Msi and epithelial markers like ECAD (**Figure 2E**). By contrast, the top 20 most mesenchymal tumors expressed lower levels of Msi and higher levels of mesenchymal markers like Fibronectin and Vimentin (**Figure 2E**). Thus, Msi expression is enriched in epithelial tumors in GBM as well, consistent with the results obtained in breast cancer tumors and cell lines.

Taken together, these results show that Msi genes are rarely mutated but frequently overexpressed across human cancers, and are strong markers of the epithelial-luminal state. This suggests that Msi proteins may play a role in the maintenance of an epithelial state and/or repression of EMT, in both breast and neural cell types. This prompted us to further explore the molecular effects of these proteins in a controlled cell culture system.

### Genetic system for inducible overexpression and depletion of Msi1/2 in NSCs

Given the upregulation of Msi genes in glioblastoma, we chose to study the molecular roles of Msi proteins in NSCs, where both proteins are highly expressed in normal development, and where their target mRNAs are likely to present. NSCs provide a well-characterized system for homogeneous cell culture (Kim et al., 2003), which is not always available for progenitor/stem cell types cultured from other primary tissues like the mammary gland, making NSCs grown in culture amenable to analysis by genome-wide techniques. Furthermore, the conserved expression of Msi genes in the nervous system and their reactivation in human glioblastoma suggests that molecular insights obtained in this system could be informative about the roles of Msi proteins in cancer cells.

We cultured cortical NSCs from E12.5 embryos obtained from transgenic mice with a Dox-inducible Msi1 or Msi2 allele, and from double conditional knockout mice for Msi1/Msi2 whose deletion was driven by a Tamoxifen (4-OHT) inducible Cre (**Figure 3A**). These systems enabled robust overexpression or depletion of Msi proteins (**Figure 3B**) within 48-72 hours of induction. To study the effects of Msi loss and gain of function on mRNA expression, processing and translation, we used ribosome footprint profiling (Ribo-Seq) (Ingolia et al., 2009) and high-throughput sequencing of polyA-selected RNA (RNA-Seq) (Mortazavi et al., 2008) (**Figure 3A**).

**Figure 3.**
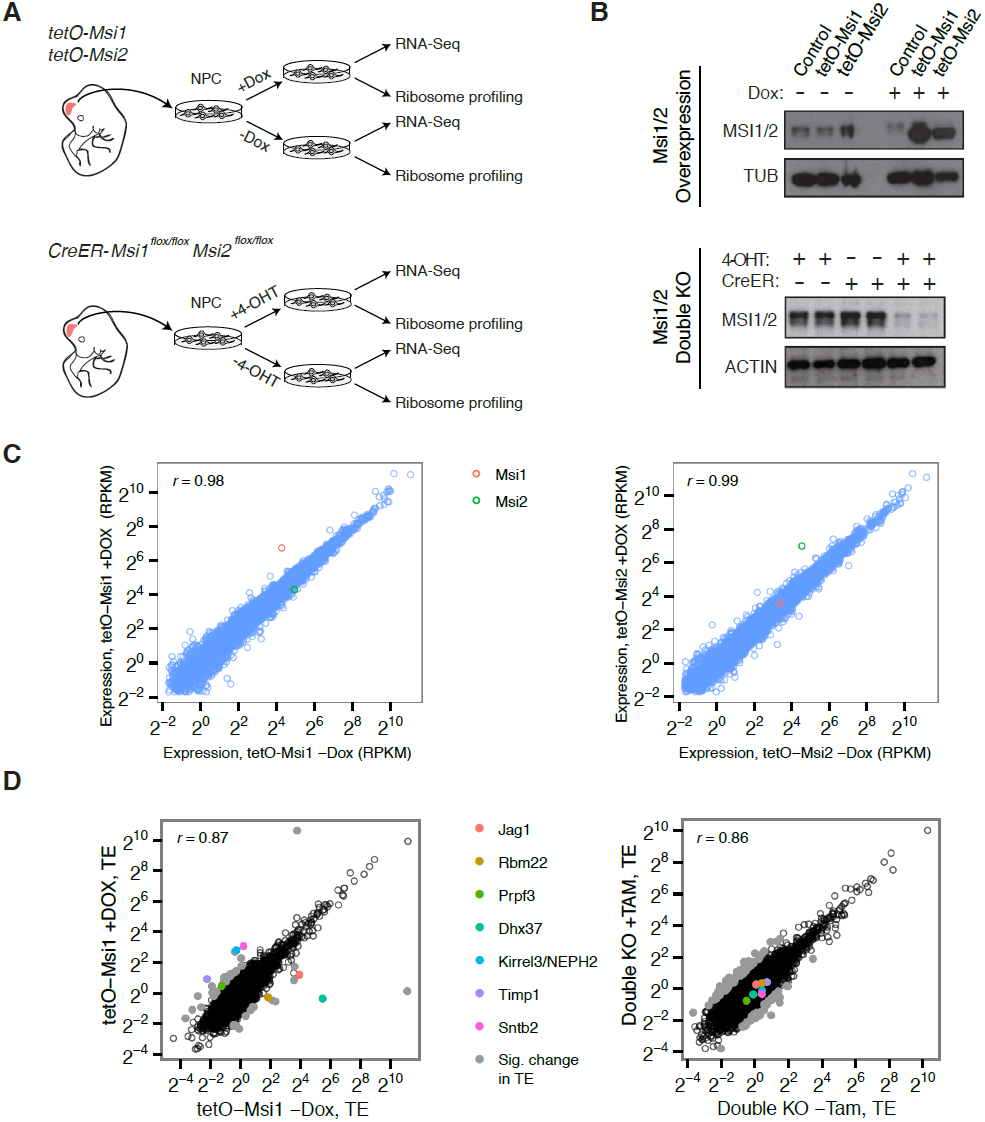
Genetic system for studying effects of Msi loss/gain of function on gene expression. **(A)** Experimental setup and use of Msi1/2 inducible overexpression and conditional double knockout mice for derivation of neural stem cells, which were then used for ribosome profiling (Ribo-Seq) and mRNA sequencing (RNA-Seq). **(B)** Western blot analysis of Musashi overexpression and knockout in neural stem cells. Overexpression and conditional knockout cells were exposed to Dox and 4-OHT for 72 hours, respectively. **(C)** mRNA-Seq expression values (RPKM) scatters between Msi1 overexpressing cells and controls (left), Msi2 overexpressing cells and controls right (72 hr Dox). Msi1/2 each robustly overexpressed with similar magnitude following Dox. **(D)** Comparison of translational efficiency (TE) values using Ribo-Seq on Msi1 overexpressing cells on Dox (72 hrs) versus controls (left) and conditional knockout cells following 4-OHT for 48 hours (right). Colored points indicate select genes with large changes in TE.

### Overexpression of Msi1 alters translation of targets without causing large changes in mRNA levels

When Msi1 or Msi2 were overexpressed, few significant changes in mRNA expression were observed after 48 hours (**Figure 3C**). This observation suggests that the regulatory activity of these factors occurs at levels other than transcription or mRNA stability/decay, such as mRNA translation. To determine the genome-wide effects of Msi proteins on translation, we performed Ribo-Seq on Msi1 overexpressing cells and double knockout cells. Ribo-Seq libraries in these experiments showed the expected enrichment for open reading frames (ORFs) over non-coding genic regions, and yielded high scores in various quality control metrics (**Supp. Figure 3**). Ribo-Seq reads were depleted from introns, and strongly enriched coding exons relative to UTRs. These quality control metrics were highly consistent across libraries, suggesting that the resulting data are comparable (**Supp. Figure 3**). To examine changes in translation, we computed Translational Efficiency (TE) values for all protein-coding genes, defined as the ratio of the ribosome footprint read density in the ORF to the RNA-seq read density, which measures ribosome occupancy along messages. Examination of TEs across overexpression and knockout samples yielded a handful of genes with very large changes in translational efficiency (**Figure 3D**).

### Msi1 represses translation of the Notch ligand Jagged1 and regulates translation of RBPs

Several genes exhibited substantial changes in their translation efficiency in response to overexpression of Msi1, including 6 with increased TE and 3 with reduced TE (**Figure 3D**). Genes with increased translation included the RNA processing factor Prpf3/Prp3p, a U4/U6 snRNP-associated factor, and genes involved in epithelial cell biology such as Kirrel3/NEPH2. Genes with repressed translation included: Rbm22/Cwc2, another splicing factor associated with U6snRNP; Dhx37, an RNA helicase; and Jag1, a ligand to Notch receptors and important regulator of Notch signaling. In order to gain insight into whether these changes are mediated by direct protein binding to RNA targets, we set out to map the sequence preferences and specificities of Msi proteins.

### Msi1 shows high affinity for RNAs containing UAG motifs

To determine the binding preferences of Msi proteins for RNA, we used “RNA Bind-n-Seq” (RBNS), a recently developed method which uses a deep sequencing approach to obtain quantitative and unbiased measurement of the spectrum of RNA motifs bound by recombinant protein *in vitro* (Lambert et al., 2014) (**Figure 4A**). Occurrence frequencies of 6mers were calculated in libraries derived from MSI1-bound RNAs and compared to their corresponding frequencies in the input library. Enrichment of 6mers was defined as the maximum fold enrichment relative to the random library across all protein concentrations. The fold enrichment profiles obtained by RBNS for the top five most enriched 6mers and five randomly chosen 6mers are shown in **Figure 4B**. Enriched 6mers exhibited similar enrichment profiles across concentrations, peaking in fold enrichment typically between 16-64 nM concentrations (**Figure 4B**). The canonical MSI1 binding site according to previous SELEX study (Imai et al., 2001) was ∼3-fold enriched by RBNS, along with highly similar sequences, showing that Bind-n-Seq can recapitulate the known binding preference of MSI1 (Ray et al., 2013). To summarize the binding preferences of MSI1 from RBNS, we aligned the top enriched 6mers and compiled them into a motif, which revealed that MSI1 binds predominantly to UAG-containing sequences (**Figure 4C**).

**Figure 4.**
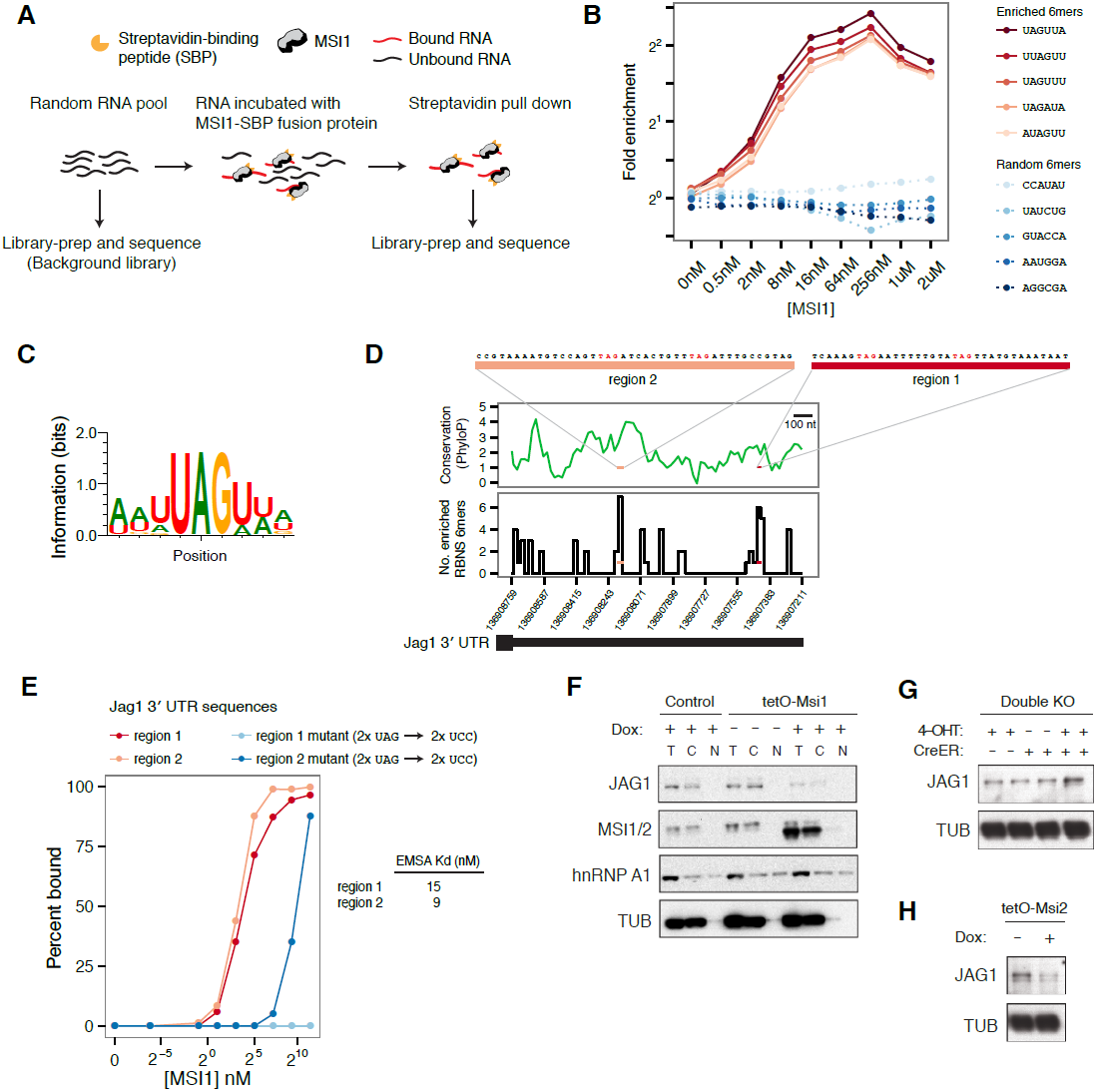
Profiling MSI1 binding preferences by RNA Bind-n-Seq. **(A)** Schemaic of Bind-n-Seq experiment for MSI1 protein. Increased concentrations of MSI1-SBP fusion protein incubated with random RNA pool, pulled by straptavidin pull-down, reverse-transcribed and sequenced. **(B)** Fold enrichment of top five enriched 6mers (red curves) and five randomly chosen 6mers (blue curves) across protein concentrations. **(C**) Binding motif for MSI1. Position-weight matrix generated by global alignment of top 20 enriched 6mers. **(D)** Two sites in Jag1 3’ UTR, region 1 and region 2, containing a high density of enriched 6mers. Top: PhyloP conservation score for 3’ UTR in 20nt windows (based on UCSC vertebrates multiple alignment). Bottom: Number of enriched 6mers from BNS in 20nt windows of 3’ UTR. (**E**) Percent binding of MSI1 protein to region 1 and region 2 (red curves) and mutants where UAG sites are disrupted (blue curves), measured by gel-shift (see **Supp. Figure 4**). Kd estimates for region 1 and region 2 are shown (mean of 2 gel-shifts per sequence). **(F)** Western blot analysis of Jag1 regulation by Msi: top left panel, Jag1 expression in Msi1 overexpression cells and controls in cellular fractions (T – total lysate, C – cytoplasmic and N – nuclear fractions). Jag1 is translationally repressed upon induction of Msi1 and detected only in total and cytoplasmic lysates. hnRNP A1, known to shuttle between the nucleus and the cytoplasm and alpha-Tubulin used as loading controls. (**G**) Increased Jag1 protein levels in double knockout cells. (**H**) Reduced Jag1 protein levels upon Msi2 overexpression.

Previous studies suggested that MSI1 binds 3’ UTR regions of mRNAs, where it acts to regulate translation (Okano et al., 2005), suggesting that the changes in translation observed in Ribo-Seq might be caused by direct binding of Msi proteins to the 3’ of these messages. To test this hypothesis in our system and validate the binding profile of RBNS, we screened the Jag1 gene (**Figure 3D**), which is translationally repressed by Msi, for occurrences of RBNS-enriched motifs. We found that the 3’ UTR of Jag1 contains a high density of RBNS-enriched 6mers (**Figure 4D**). We selected two regions of the Jag1 3’ UTR that contained the highest density of RBNS-enriched 6mers in order to test whether these sequences can bind the MSI1 protein *in vitro* (**Figure 4B**, top). Both region sequences were found to bind tightly to MSI1 by gel-shift assay (region 1: 15 Kd, region 2: 9 Kd – see **Supp** **Figure 4D** for representative gel shifts). Since both sequences contain UAG motifs (**Supp. Figure 4D**), we hypothesized that the UAG sites are required for binding. Consistent with this, mutation of the UAG sites to UCC in both sequences either fully or near-fully ablated reduced binding to MSI1 (**Figure 4E**).

Following Msi overexpression, Jag1 had ∼5-fold lower Ribo-Seq while its mRNA level was little changed, suggesting a predominant effect at the translational level (**Supp Figure 5**). In double knockout cells, Jag1 mRNA levels were upregulated ∼1.5 fold (**Supp Figure 5**), as measured both by RNA-Seq and Ribo-Seq, suggesting effects on message stability either in the absence of or as a consequence of translational derepression. Western blot analysis confirmed the repression of Jag1 by Msi1 (**Figure 4F**) and its upregulation in double knockout cells (**Figure 4G**). Given the high degree of homology between Msi1 and Msi2 at the protein level, we predicted that Msi2 overexpression would similarly repress Jag1, and this was confirmed by western blot analysis (**Figure 4H**). These results provide functional validation for the RBNS-derived binding site, and support a model where Msi proteins directly bind to the Jag1 3’ UTR in order to regulate translation or message stability.

**Figure 5.**
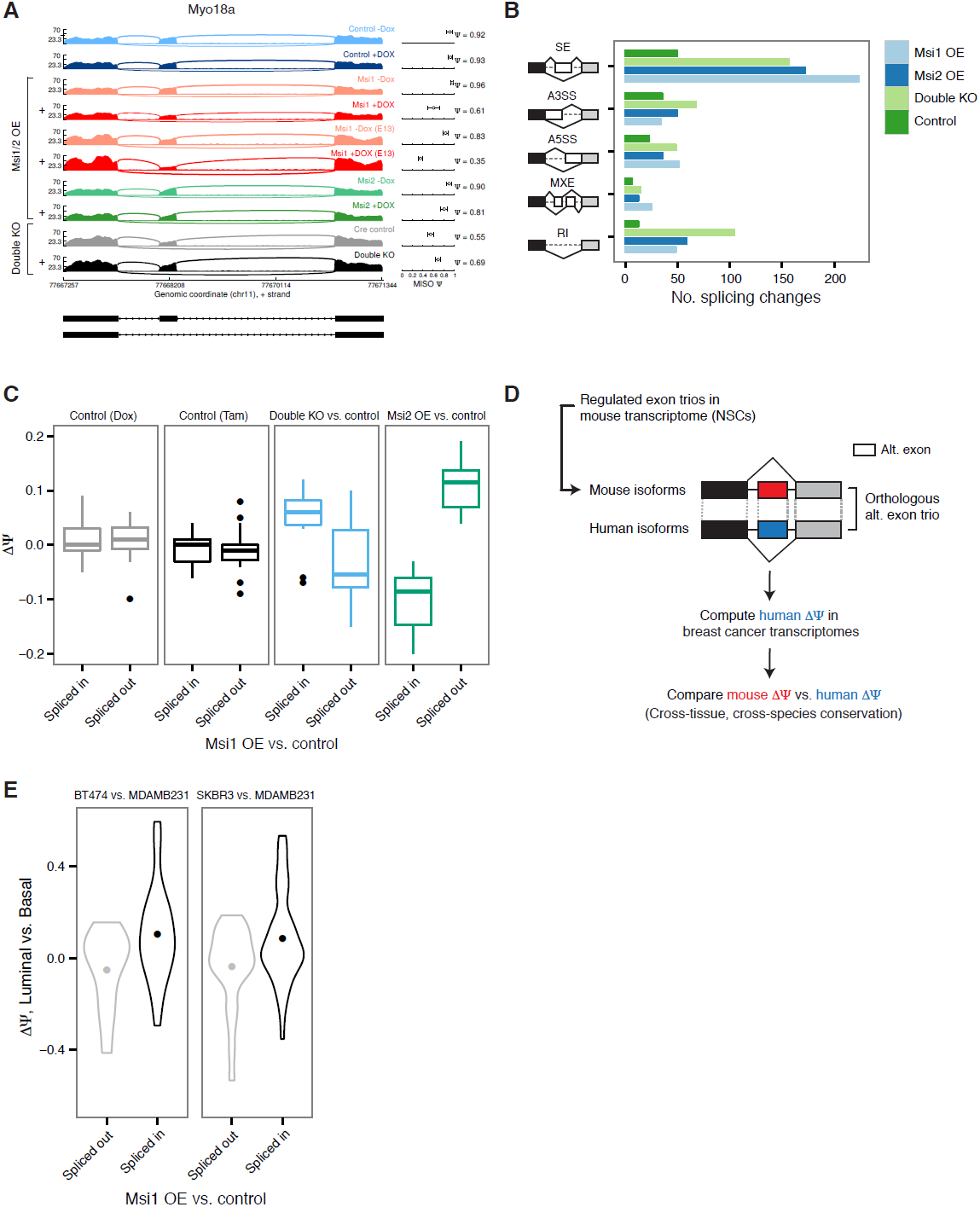
Global impact of Msi proteins on alternative splicing. **(A)** Sashimi plot for Myo18a alternative exon 38 with Percent Spliced In (Ψ-value) estimates by MISO (values with 95% confidence intervals, right panel.) Exon splicing is repressed by Msi1 overexpression and slightly increased in knockout Msi1/2 cells. ‘**+**’ indicates samples treated with Dox/Tam for overexpression/knockout cells, respectively. E12.5 neural stem cells were used for all samples except Msi1 overexpression for which an additional E13.5 NPC time point was sequenced. **(B)** Number of differential events (MISO Bayes factor ≥ 10, ΔΨ ≥ 0.12) in each alternative RNA processing category (SE – skipped exons, A5SS – alternative 5’ splice site, A3SS – alternative 3’ splice site, MXE – mutually exclusive exons, RI – retained introns) for Msi1 overexpression (‘Msi1 OE’), Msi2 overexpression (‘Msi2 OE’), double knockouts (‘Double KO’), and a Dox control pair (‘Control’). **(C)** Comparison of ΔΨ in Msi1 overexpression versus control binned by direction (‘Spliced in’ or ‘Spliced out’, x-axis) to ΔΨ in Msi2 overexpression cells and in double knockout cells (along with respective Tam and Dox controls, y-axis). **(D)** Computational strategy for identifying human orthologs of alternative exon trios regulated in mouse neural stem cells. Orthologous exon trios were identified by synteny using multiple genome alignments. **(E)** Comparison of ΔΨ mouse alternative exons by Msi1 (comparing overexpression to control, x-axis) and ΔΨ of their orthologous exon trios in human (comparing luminal and basal cell lines, y-axis). Two pairs of luminal and basal cells compared: BT474 vs. MDAMB231 and SKBR3 vs. MDAMB231. ΔΨ value distributions summarized by violin plots with a dot indicating the mean ΔΨ value.

### Indirect regulation of alternative splicing by Msi proteins

Since some of the largest changes in translation observed by Ribo-Seq affected splicing-associated RBPs, we hypothesized that Msi overexpression might alter pre-mRNA splicing globally. We assessed changes in mRNA splicing following Msi overexpression or depletion in RNA-Seq data using the MISO software (Katz et al., 2010). For example, exon 38 in the Myo18a gene, which is predominantly included under control conditions, is modestly repressed following Msi2 overexpression and strongly repressed following overexpression of Msi1 (**Figure 5A**). Conversely, exon 21 in Erbin (Erbb2ip; a direct binding-partner of the breast cancer oncogene HER2/Erbb2) is strongly enhanced by Msi1 overexpresssion (**Figure 5B**). Overall, we observed several hundred differentially spliced exons that were either repressed or enhanced by overexpression/knockout of Msi proteins (**Figure 5C**). The predominant localization of Msi proteins is to the cytoplasm (**Supp. Figure 6**) even when overexpressed (**Figure 3F**), suggesting that these changes in pre-mRNA splicing are indirect, e.g., they may result from changes in the levels of splicing factors resulting from translational regulation of their mRNAs by Msi.

**Figure 6.**
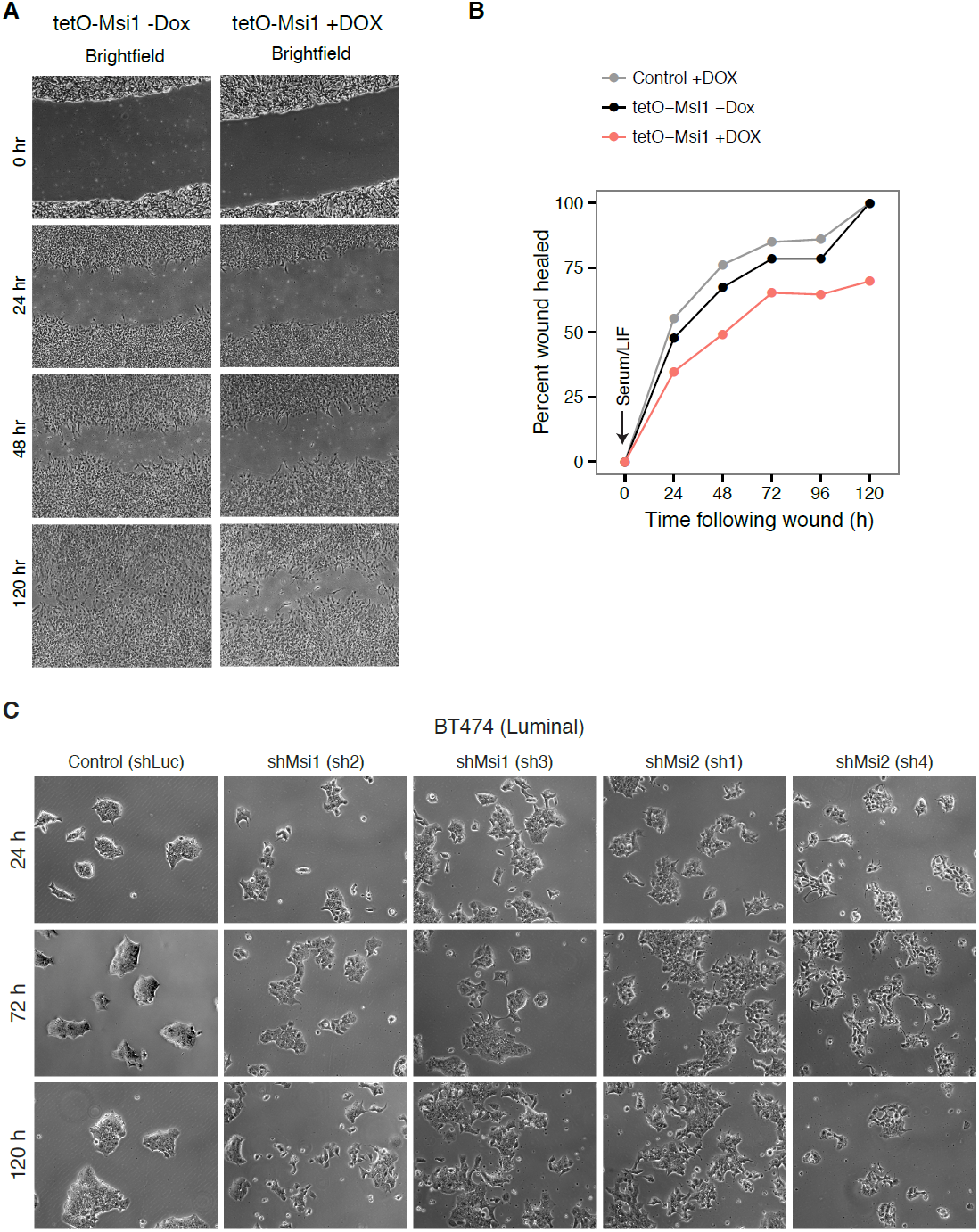
Msi levels alter EMT processes and epithelial morphology in mouse NSCs and breast cancer cell lines. **(A)** Wound healing assayed in tetO-Msi1 cells on Dox for 24-48 hours and then scratched in LIF/Serum medium, which induces EMT in embryonic neural stem cells. Left panel: control cells. Right panel: Msi1 overexpressing cells. **(B)** Automated quantification of the percentage of wound healed across time in Msi1 overexpressing cells (red) and Dox-free control cells (black) and control cells with Dox (grey). (**C**) Knockdown of Msi-1/2 in BT474 breast cancer cell line using lentiviruses carrying short hairpins (shRNAs). Brightfield images (10x magnification) shown at 24, 72, and 120 hours after Puromycin-selection.

### Msi1 and Msi2 cause similar global effects on mRNA splicing

Mouse MSI1 and MSI2 proteins are over 70% identical at the amino acid level and contain highly similar RNA recognition motifs, suggesting that they may be at least partially functionally redundant. To test whether Msi1 and Msi2 exert similar effects on mRNA splicing, we correlated the observed direction of splicing changes following Msi1 or Msi2 overexpression. Exons with increased inclusion following Msi1 overexpression tended to show increased splicing in Msi2 overexpression conditions as well, and similarly for those with decreased inclusion (**Figure 5D**). These observations suggested that Msi1 and Msi2 exert similar effects on mRNA splicing when overexpressed. The pattern of Msi1 overexpression-induced splicing changes was uncorrelated with Dox-induced splicing changes in control cells, arguing against an effect of Dox alone. Comparing the directions of Msi1-induced splicing changes with those observed in double knockout cells exposed to 4-OHT (**Figure 5D**), we observed that Msi1 overexpression induced changes that negatively correlated with splicing changes seen in the Msi1/2 double knockout cells (**Figure 5D**). This observation supports that the observed effects are part of the normal function of Msi proteins rather than an artifact of Msi overexpression. We observed no correlation in splicing between Msi1-induced splicing changes with those seen in the control condition of double floxed cells exposed to 4-OHT but lacking the Cre driver (**Figure 5D**).

### Splicing program induced by Msi in stem cells is conserved in human cancer cell lines and associated with luminal state

Our observation that splicing is altered when Msi expression is perturbed in mouse NSCs raises the question of whether this function is conserved in human breast cancer cells, and whether the program might be associated with a particular cell state. The natural variation in Msi levels across breast cancer cell lines (**Figure 2C-E**) enables a comparison of cancer transcriptome splicing patterns between Msi-high versus Msi-low cell types. We next used this variation in Msi levels to gain insight into whether the Msi induced splicing program in mouse is conserved in human breast cancer cell lines.

If splicing patterns induced by Msi were conserved from mouse to human, one would expect splicing changes in human samples with high versus low Msi levels to match the direction of change in mouse Msi overexpression. To test this, we identified orthologous human alternative exon trio for each mouse alternative exon using synteny in a multiple genome alignment (**Figure 7A** **and Supp. Methods**). Using these homologous exon trios, we asked whether the pattern of splicing induced by Msi in murine cells is recapitulated in human samples. We calculated the Ψ values of human alternative exon trios in breast cancer cell lines (**Supp. Methods**). We first compared ΔΨ for alternative exons regulated by Msi in mouse, between Msi1 overexpressing cells and controls, to ΔΨ values of their orthologous trios between luminal and basal breast cancer cell lines (**Figure 7B**). The splicing patterns were correlated: exons spliced in upon Msi1 overexpression in mouse had higher Ψ values in luminal (Msi-high) than in basal (Msi-low) cell lines, and vice versa for exons spliced out upon Msi1 overexpression in mouse (**Figure 7B**). This correlation was observed in several different breast cancer luminal and basal pairs, but was strongest when comparing HER2+ luminal lines such as BT474 and SKBR3 to basal lines, consistent with the observation that Msi levels are higher in these HER2+ cell lines (**Figure 2D**). This correspondence suggests that the Msi induced splicing changes are conserved from mouse to human and across cell types, and that the induced splicing pattern matches that of the epithelial-luminal state.

Two of the most strongly affected alternative exons in murine NSCs, in the Myo18a and Erbin genes (**Figure 5A-B**), were conserved in the human genome and detected in the transcriptomes of all breast tumors and controls. In primary tumors, these events showed a striking cancer-associated splicing pattern, with Erbin exon 21 splicing enhanced in tumors and Myo18a exon 38 splicing repressed in tumors **(Supp.** **Figure 7A**). A model where Msi proteins regulate these alternatively spliced exons in tumors would predict that the extent of Msi overexpression in a tumor would correlate with the magnitude of the effect on exon splicing. To test whether the regulation of these exons is responsive to Msi levels, we correlated the fold change in Msi expression for each matched tumor-control pair with the ΔΨ value of the Erbin and Myo18a exons in that pair (**Supp. Figure 7B**). We observed high correlation (particularly for Msi2) between the extent of Msi overexpression and the change in splicing in luminal tumors. As in mouse neural stem cells, increased expression of Msis was associated with increased inclusion of the Erbin exon and repression of Myo18a exon splicing, suggesting that Msi-dependent regulation of splicing is conserved in primary tumors in addition to breast cancer cell lines.

### Msi overexpression inhibits EMT processes in murine NSCs

The Notch pathway regulator Jag1, which we found was translationally repressed by Msi, is known to be required for EMT. Jag1-depleted keratinocytes undergoing TGFβ-induced EMT fail to express mesenchymal markers and retain epithelial morphology (Zavadil et al., 2004). Furthermore, knockdown of Jag1 in keratinocytes strongly impairs wound healing (Chigurupati et al., 2007), a process that requires cells to acquire mesenchymal properties such as migration and protrusion. Our gene expression analysis also further supported the mesenchymal-basal specific expression of Jag1, which is particularly pronounced in breast cancer (**Figure 2**). The epithelial-associated expression pattern of Msi genes and the antagonistic relation between Msi and Jag1 (**Figure 2**) prompted the hypothesis that Msi activation promotes an epithelial cell identity, effectively blocking EMT.

To test the hypothesis that Msi activation may promote an epithelial state and hinder EMT, we assessed the effect of Msi overexpression on wound healing. Embryonic NSCs cultured with LIF/Serum have been observed to undergo an EMT (Ber et al., 2012). We found that when Msi1 was overexpressed cells were severely impaired in wound healing following stimulation with LIF/Serum (EMT medium) prior to wounding (**Figure 6A-B**). Exposure to LIF/Serum acutely blocks proliferation of NSCs and overexpression of Msi1/Msi2 in the absence of LIF/Serum did not alter proliferation rates (data not shown), ruling out differences in proliferation as a cause the differences in wound healing. Control cells on Dox were not impaired in this process (**Figure 6A-B**), and NSCs in standard non-EMT medium overexpressing Msi1 or Msi2 were also impaired in wound healing while double knockout cells showed modest acceleration in would healing (data not shown).

### Msi proteins are required for maintenance of the epithelial-luminal state in breast cancer cells

To address whether Msi proteins are functionally required for the maintenance of the luminal state, we knocked down Msi1/Msi2 in two luminal breast cancer cell lines (BT474, MCF7-Ras) where Msi proteins are highly expressed (**Figure 6C**, **Supp Figure 8A**). In the HER2+ luminal cell line BT474, cells grow in tightly packed epithelial colonies. We observed a striking morphological change upon knockdown of Msi1 or Msi2, where cells progressively separated and acquired a basal-like appearance with 3-5 days of knockdown (**Figure 6C**), accompanied by reduced proliferation (data not shown). A similar phenotype was observed in MCF7-Ras cells upon knockdown of Msi1 or Msi2 (**Supp. Figure 8B**). These results argue that Msi expression is required for the maintenance of the epithelial-luminal state in breast cancer cell lines.

### Msi2 overexpression in basal cell layer of mammary gland results in defective and delayed ductal branching morphogenesis

In light of Msis association with the luminal state in breast cancer tumors and their effect on the epithelial-luminal state in breast cancer cell lines, we next asked whether Msi proteins play similar roles in the mammary gland *in vivo*. During maturation epithelial cells in the mammary gland migrate and form ducts within the mammary fat pad through a process termed mammary ductal branching morphogenesis. The formation of the mammary ductal system is thought to be a kind of EMT, making mammary gland an attractive system to study the regulation the transition *in vivo*.

The mammary gland Terminal End Buds (TEBs) from which ducts form is organized into discrete layers of cell types, including epithelial luminal and basal cells. The identity of luminal and basal tumors is thought to resemble the their mammary gland cell type counterparts. mRNA-Seq expression analysis of purified mouse mammary luminal (CD24^high^CD29^+^) and basal (CD24^+^CD29^high^) cells from (dos Santos et al., 2013) revealed enrichment of Msi1 and Msi2 expression in luminal cells (data not shown.) As predicted by the mRNA expression profile, Msi2 protein level was highest in the luminal cell layer and far lower in the basal (K14-positive) cell layer (**Figure 7A**).

**Figure 7.**
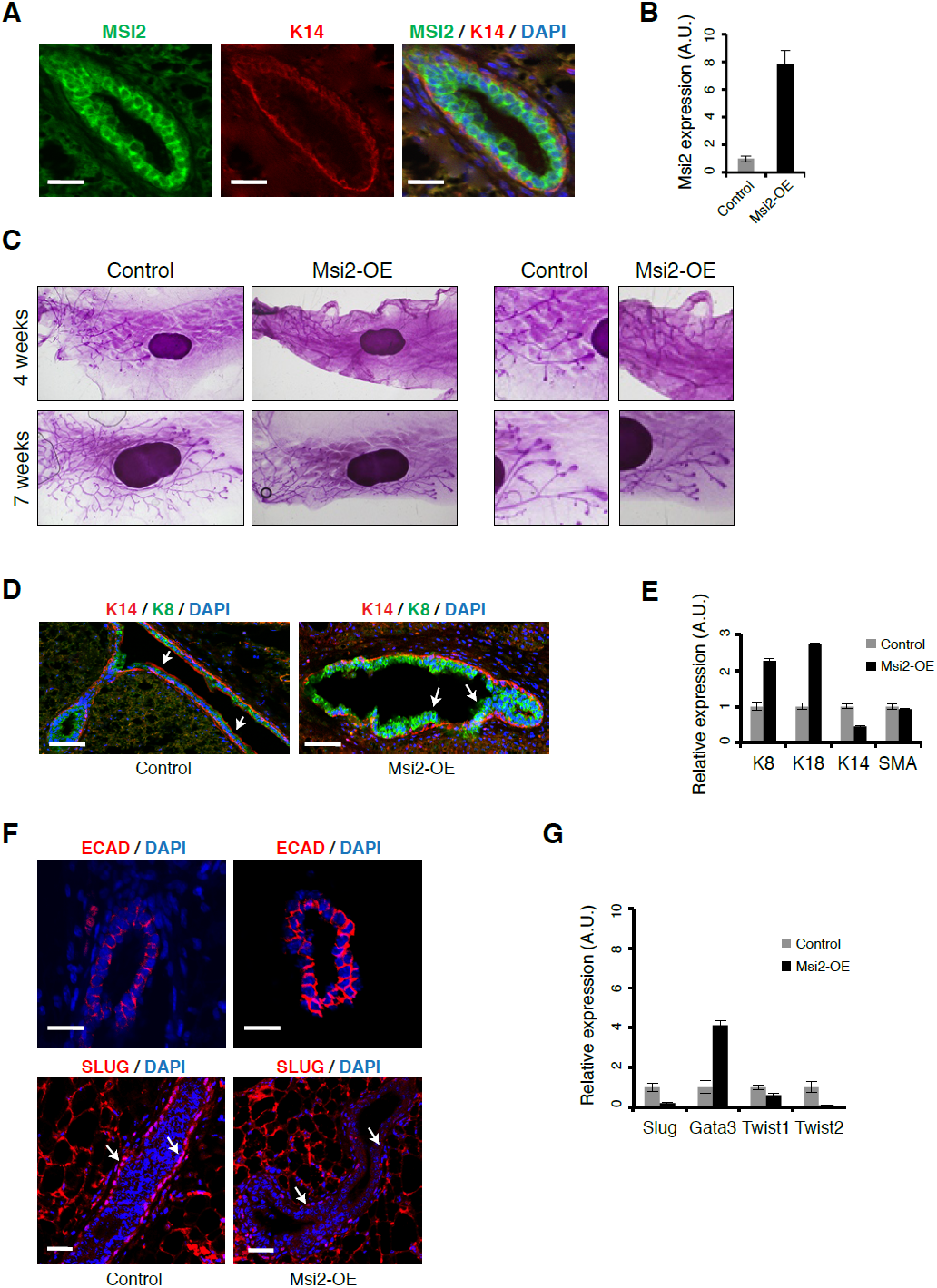
Msi2 activation represses EMT and expands mammary luminal cell layer *in vivo*. **(A)** Immunostaining for MSI2, K14 and DAPI in control sections of mammary gland. Scale bar: 50 μm **(B)** qRT-PCR for Msi2 in mammary epithelial cells from control and Msi2 overexpressing mice (“Msi2-OE”). **(C)** Whole mount stain for mammary glands from control and Msi2 overexpressing mice (left: low magnification, right: high magnification.) **(D)** Immunostaining for K14, K8 and DAPI in mammary gland sections from control and Msi2 overexpressing mice. Scale bar: 100 μm **(E)** qRT-PCR for luminal markers (K8, K18), basal markers (K14), and smooth-muscle Actin (SMA) in mammary epithelial cells from control and Msi2 overexpressing mice. **(F)** Staining for E-cadherin (ECAD) (top) and EMT-marker SLUG (bottom) in mammary glands from control and Msi2 overexpressing mice. Luminal cell layer is expanded upon Dox (arrows). Scale bar: 100 μm. **(G)** qRT-PCR for Slug, Gata3, Twist1, Twist2 in mammary epithelial cells from control and Msi2 overexpressing mice. Slug expression in basal cell layer is reduced upon Dox (arrows). Scale bar: 50 μm.

#### Msi2 overexpression drives expansion of luminal cell layer and blocks EMT *in vivo*

We next wanted to investigate the effect of Msi overexpression on epithelial cell state in the mammary gland in order to see whether its *in vivo* effects on epithelial-luminal state are similar to those observed in culture models. We ectopically expressed Msi2 in the basal cell layer, where it is nearly absent normally (**Figure 7A**), using a basal cell-specific Dox-inducible driver, K14-rTTA. As expected, mice administered Dox showed significantly higher levels of Msi2 protein in the basal cell layer (**Supp. Figure 9A**) and overall higher levels of Msi2 mRNA in mammary epithelial cells (**Figure 7B**). Overexpression of Msi2 resulted in a defective and delayed mammary ductal branching pattern (**Figure 7C**). Since branching morphogenesis requires cells to lose their epithelial identity and undergo migration, we hypothesized that the defect in branching morphology might be due to the inability of cells to lose their epithelial identity and/or due to the expansion of an epithelial cell layer.

Consistent with this hypothesis, we observed that Msi2 overexpression resulted in expansion of the luminal cell layer **(Figure 7D, Supp. Figure 9B)**. We found that luminal cell marker expression increased while basal marker expression decreased (**Figure 7E**), reflecting the change in cell type distribution. These results support a model where Msi ectopic expression leads to expansion of epithelial-luminal cells in the mammary gland, effectively blocking EMT processes needed to undergo branching morphogenesis, and resulting in the defective ductal elongation observed in **Figure 7C**. To directly test this hypothesis, we examined the expression of EMT markers upon Msi2 overexpression. Msi2 overexpression led to an increase in epithelial marker E-cadherin and reduction in Slug, a marker of EMT and mesenchymal cells. mRNA levels of the Slug and the EMT/mesenchymal regulators Twist1 and Twist2 decreased upon Msi2 overexpression, while the marker of luminal epithelial cells Gata3 increased (**Figure 7G, Supp. Figure 9C**), consistent with a model where Msi2-induced expansion of the luminal cell layer blocks EMT. In sum, Msi2 functions as a regulator of epithelial cell state in the mammary gland that blocks EMT when ectopically expressed, mirroring the functions of Msi proteins that we observed in breast cancer cell lines and neural stem cells. These results suggest that Msi proteins play a similar role in a healthy *in vivo* context in the mammary gland to that seen in cancer cells.

## Discussion

### Post-transcriptional control of cell state by RBPs

Our data show that Msi proteins regulate mRNA translation and splicing, the latter likely through regulation of splicing factors, and that their activation promotes an epithelial cell state (**Figure 8**). The commonalities between our data in mouse and human cell types additionally suggest that these functions are conserved across species. Our work contributes to a growing body of evidence that RBPs might be comparably important to transcription factors in the control of cell states. The varied localization patterns of RBPs within the cell allow for subtle changes in mRNA processing and translation, and suggest that merely profiling mRNA levels is not sufficient to determine the effects of these proteins on cell state.

**Figure 8.**
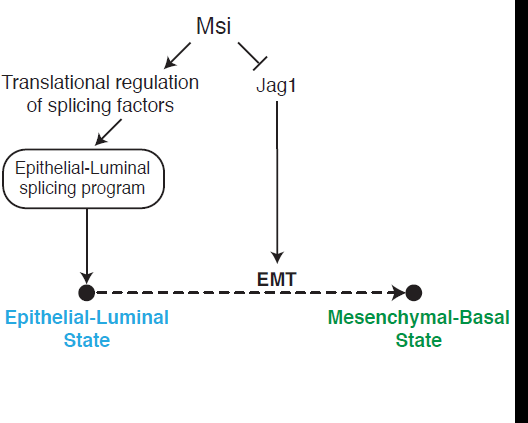
Model for Msi roles in regulation of cell state. Model for Msi role in the control of the epithelial state.

Like the Msi family, the RBPs Esrp1/Esrp2 are also enriched in epithelial cells, but are localized to the nucleus where they directly regulate splicing (Warzecha et al., 2009). ESRPs promote an epithelial splicing program that is eliminated during EMT. Ectopic expression of Esrp1 alone can induce epithelial features in mesenchymal cells, highlighting the importance of RBPs as drivers of cell state transitions that are central to cancer cells (Reinke et al., 2012). Recently, it was proposed that Snail acts to promote the mesenchymal state in part by repressing Esrp1 and that ectopic expression of Esrp1 alone can induce epithelial features in mesenchymal cells, further highlighting the importance of RBPs as epithelial state regulators (Reinke et al., 2012). Our results show that Msi proteins are cytoplasmic-localized analogs of the Esrp family, exerting their effects primarily through regulation of cytoplasmic RNA rather than direct regulation of splicing in the nucleus. The molecular mechanism by which Msi proteins exert specific translational effects is not fully understood, through a model where these proteins repress translation by outcompeting the RNA helicase eIF4G for PolyA-binding protein (PABP) was proposed (Kawahara et al., 2008).

Similarly to Esrps, the nuclear splicing factor SF2/ASF plays a major role in the cancer transcriptome through post-transcriptional regulation of tumor suppressor splicing and is oncogenic when overexpressed (Karni et al., 2007). Initial studies comparing epithelial and mesenchymal cancer transcriptomes found several more RBPs enriched in the epithelial cancer state (Shapiro et al., 2011), suggesting that RBPs play a broader role in the maintenance of this state and can be used to manipulate cell state transitions in cancer.

### Roles of Jagged1 and Notch signaling in breast cancer tumors and EMT

The role of Notch signaling in cancer remains complex and appears to vary between cancer types and subtypes (Dickson et al., 2007; Lobry et al., 2011). The upregulation of Jag1 in the basal state suggests that Notch pathway activity is high in and required for entry into the mesenchymal state, consistent with previous studies (Dickson et al., 2007; Zavadil et al., 2004). In mammary epithelial cells, Jag1-triggered activation of Notch was shown to reduce Ecadherin expression and increase Slug expression (Leong et al., 2007). Furthermore, Jag1 activation in breast cancer cells promotes their metastasis into the bone *in vivo* by activating Notch in neighboring bone cells (Sethi et al., 2011). The dependence of EMT on Notch activation has been observed in normal development as well. During heart development, cardiac valves are generated from endocardium through EMT, and Notch activity was shown to be required for this process (Timmerman et al., 2004). Collectively, these studies are consistent with our working model in which Msi repress Jag1 translationally, in turn altering Notch activity required for EMT, in addition to other pleiotropic effects Jag1 may have apart from its role as Notch ligand. Our findings reveal an additional layer of translational control of Jag1 levels and it would be interesting to explore the spatial, non-cell-autonomous effects of Jag1 repression in mammary glands and mammary tumors.

### Broad regulation of the epithelial stem/progenitor cell state

Msi proteins are co-expressed with various proliferation markers in a wide variety of stem cell niches, including the breast, stomach, intestine, lung and brain, leading to the hypothesis that these are general epithelial stem cell/progenitor regulators across tissues. Our findings are consistent with this hypothesis, though uncertainty surrounding the identity of stem cell types in adult tissues makes it difficult to test directly. Our work suggests that would be fruitful to pursue the role of Msi in the normal development and transformation of other adult tissues. The lung, like the mammary gland, is a relevant system for studying Msi overexpression in light of our observation that Msi is frequently overexpressed in lung tumors. Finally, the systematic downregulation of Msi1/Msi2 and high frequency of Msi1 mutations in kidney tumors suggests that the kidney would be an informative model for studying Msi loss-of-function and its consequences in cancer.

## Author contributions

YK conceived project, performed majority of experiments, all computational analyses, and wrote manuscript. FL and ZY performed *in vivo* mammary gland experiments. NL performed RBNS and associated EMSA analysis. ES, WT, AWC, and CJL contributed reagents and expert advice. EMA, PBG contributed to data analysis and contributed expert advice. RJ and CBB supervised project and contributed to writing of manuscript and interpretation of results.

## Methods

### Mouse strains and derivation of neural stem cell lines

Mice of the 129SvJae strain were used. For derivation of embryonic neural stem cells (NSCs), littermate embryos were used whenever possible. Cortical NSCs were derived from embryos following (Kim et al., 2003). Briefly, cortical tissue was isolated from E12.5 embryos (unless otherwise noted) under a light dissection microscope inside a sterile fume hood and collected by centrifugation. Cortical tissues were dissociated into single cells by trituration in Magnesium/Calcium-free HBSS buffer (Gibco) followed by 15 min incubation at room temperature. Dissociated tissue was collected by centrifugation, resuspended in N2 medium containing growth factors and Laminin (Life Technologies), and plated onto Polyornithin/Laminin-coated tissue culture dishes as in (Okabe et al., 1996).

### Culture conditions for embryonic neural stem cells

NSCs were grown in N2 medium (Okabe et al., 1996) containing EGF (20 ng/ml) and bFGF (20 ng/ml) and Laminin (Life Technologies). Cells were grown on Polyornithin/Laminin-coated dishes. EMT was induced by switching cells to N2 medium containing LIF/FBS as described in (Ber et al., 2012).

### Culture conditions for human breast cancer lines and shRNA knockdowns

All breast cancer lines were cultured in DME containing 10% FBS, 1% GlutaMAX (Gibco), and Penn/Strep, except for BT474, which was cultured in RPMI base medium, and SKBR3 which was cultured with McCoy’s 5A supplement. Lentiviruses carrying pLKO vectors with hairpins against Msi1, Msi2 or Luciferase (control) were used for knockdowns. Hairpins were obtained from Broad Institute shRNA library. Cells were infected in a centrifuge spin-infection step (1500 RPM, 37 C, 20 mins) and viral medium was left on cells overnight. Cells were subjected to 4-6 day Puromycin selection (2 ug/ml) 48 hours after infection.

### Western blotting, immunofluorescence staining and antibodies used

For western blotting, cells were lysed on ice and protein lysates were loaded onto 4-12% gradient Bis-Tris Gel (Life Technologies). Primary antibodies and dilutions used in western blotting: anti-MSI1/2 (Cell Signaling Technology #2154, 1:800), anti-MSI2 (Abcam #57341, 1:800), anti-Jag1 (Cell Signaling Technology #2620, 1:800), anti-HER2 (Cell Signaling Technologies #2248, 1:1000), anti-phos-HER2 (Cell Signaling Technology #2241, 1:1000), anti-alpha-Tubulin (Sigma-Aldrich T9026, 1:5000), anti-HNRNPA1 (Abcam ab5832, 1:800). Immunofluorescene was performed on cells grown on glass bottom chambers (LabTek II, #1.5), fixed in 4% PFA. Cells were blocked and permeabilized in 5% FBS, .1% Triton in PBS(+). Antibodies were applied in 1% FBS in PBS(+). Immunofluorescence antibodies and dilutions: anti-MSI1 (MBL D270-3, 1:500), anti-HNRNP A2/B1 (Santa Cruz, sc-374052, 1:200).

### Immunohistochemistry on human breast cancer sections

Paraffin-embedded human breast cancer sections were obtained from Biomax US (BR1505a) and stained using standard protocols with antigen retrieval. Antibodies used: anti-ECAD1 (BD Biosciences, 1:50) and anti-MSI1 (MBL D270-3, 1:200).

### Confocal imaging for immunofluorescence

Confocal imaging was performed using a Perkin-Elmer microscope using oil-immersion 63x objective, imaged with Velocity software. Single confocal stacks or maximum Z intensity projections were obtained using Fiji (Bioformats-LOCI plugin).

### RNA-Seq and ribosome profiling library generation

RNA-Seq libraries were prepared from polyA-selected RNA using standard Illumina protocol. Ribosome profiling libraries were prepared following (Ingolia et al., 2009) with several modifications. Briefly, cells were collected by centrifugation and immediately flash-frozen. Cells were thawed in lysis buffer (20 mM HEPES [pH 7.0], 100 mM KCl, 5 mM MgCl2, 0.5% Na-Deoxycholate, 0.5% NP-40, 1 mM DTT, Roche mini EDTA-free protease inhibitor tablets [1 tablet/10 ml]) and briefly treated with DNase I and RNAse I. Nuclei and cell debris were removed by centrifugation and lysates were treated with RNase I (NEB) for 75 mins at room temperature to generate monosome-protected RNA fragments. Monosomes were collected by ultracentrifugation in a sucrose cushion, denatured in 8 M Guanidium HCl, and protected RNA fragments (footprints) were extracted with Phenol-Chloroform. Footprints were dephosphorylated by PNK treatment and size-selected (∼31-35 nt fragments) by purification from a 15% TBE-Urea gel. Subtractive hybridization of ribosomal RNA from footprints was performed as in (Wang et al., 2012). Footprints were then polyA-tailed, and Illumina sequencing adaptors were added in a reverse transcription step to obtain footprint cDNA, which was then isolated by gel purification. cDNA was then circularized, PCR-amplified, and PCR products isolated by gel purification and submitted for sequencing on Illumina Hi-Seq platform.

### Computational analysis of RNA-Seq, ribosome profiling and Bind-n-Seq

Source code for the pipelines used to analyze RNA-Seq, ribosome profiling and Bind-n-Seq data is available through the open-source library <u>rnaseqlib</u> (available at the git repository: http://www.github.com/yarden/rnaseqlib). Detailed analysis procedures, gene lists and additional information about all genomic datasets are available at: http://www.musashi-genes.org

### Sequencing data availability

All RNA sequencing data was submitted to GEO (accession GSE58423).

### Computational analysis of TCGA data

Publicly available TCGA data sets (Level 2 and Level 3) were downloaded from NIH ‘Bulk Download’ website. RNA-Seq analyses were performed using ‘RNASeqV2’ TCGA files. Fold changes for genes were normalized by correction with Lowess-fit of MA-values calculated using raw gene expression estimates. Alternative exon expression was quantified using MISO.

### Computational identification of orthologous exon trios between mouse-human

Syntenic regions for exons in mouse alternative exon trios (mm9) were computed using Ensembl Compara Database (Release 66) PECAN multiple genomes alignment, using the Pycogent Python framework (Knight et al., 2007). Syntenic coordinates in human genome (hg19) were then matched to annotated hg19 exon coordinates given in TCGA data files.

### RNA Bind-n-Seq method (RBNS) Cloning and protein expression

A streptavidin binding peptide (SBP) tag was added to the pGEX6P-1 vector (GE) after the Presceission protease site. Full length Musashi (MSI1) was cloned downstream of the SBP tag with infusion (Clonetech) using BamHI and NotI cloning sites. Msi expression was induced with 0.5 mM IPTG at 18 degrees for 4 hours in the Rosetta(DE3)pLysS E. coli strain and subsequently purified on a GST GraviTrap column (GE). MSI1 was eluted from the GST column with PreScission protease (GE) in 4 mL of Protease Buffer (50 mM Tris pH 7.0, 150 mM NaCl, 1mM EDTA, 1 mM DTT) at 4 degrees overnight (∼16 hours). Protein purity was assayed SDS-PAGE gel electrophoresis and visualized with SimplyBlue SafeStain (Invitrogen).

### Random RNA preparation

Input random RNA was generated by T7 in vitro transcription. 1 μg T7 oligo was annealed to 1 μg of RBNS T7 template by heating the mixture at 65 degrees for 5 minutes then allowing the reaction to cool at room temperature for 2 minutes. The random RNA was then in vitro transcribed with HiScribe T7 In vitro transcription kit (NEB) according to manufacturers instructions. The RNA was then gel-purified from a 6% TBE-urea gel.

#### RBNS T7 template

~~~
CCTTGACACCCGAGAATTCCA(N)40GATCGTCGGACTGTAGAACTCCCTATAGTGAGTCGTATTA
~~~

#### T7 oligo

~~~
TAATACGACTCACTATAGGG
~~~

#### Resulting RNA Pool

~~~
GAGTTCTACAGTCCGACGATC(N)40TGGAATTCTCGGGTGTCAAGG
~~~

### MSI1 binding assay

Nine concentrations of purified MSI1 (0 nM, 0.5 nM, 2 nM, 8 nM, 16 nM, 64 nM, 256 nM, 1 μM and 2 μM) were equilibrated in 250 ul of Binding Buffer (25 mM Tris pH 7.5, 150 mM KCl, 3 mM MgCl2, 0.01% Tween, 1 mg/mL BSA, 1 mM DTT, 30 μg/mL poly I/C (Sigma)) for 30 minutes at room temperature. 40 U of Superasin (Ambion) and 1 μM random RNA (final concentration) was added to the MSI1 solutions and incubated for 1 hour at room temperature. During this incubation, Streptavidin magnetic beads (Invitrogen) were washed 3 times with 1 mL of wash buffer (25 mM Tris pH 7.5, 150 mM KCl, 60 μg/mL BSA, 0.5 mM EDTA, 0.01% Tween) and then equilibrated in Binding Buffer until needed. MSI1 and interacting RNA was pulled down by adding the RNA/protein solutions to 1 mg of washed streptavidin magnetic beads and incubated for one hour at room temperature. Supernatant (unbound RNA) was removed from the beads and the beads washed once with 1 mL of Wash Buffer. The beads were incubated at 70 degrees for 10 minutes in 100 μL of Elution Buffer (10 mM tris pH 7.0, 1 mM EDTA, 1% SDS) and the supernatant collected. Bound RNA was extracted from the eluate by phenol/chloroform extraction and ethanol precipitation. Half of the extracted RNA from each condition was reverse transcribed into cDNA using Superscript III (Invitrogen) according to manufacturer’s instructions using the RBNS RT primer. To control for any nucleotide biases in the input random library, 0.5 pmol of the RBNS input RNA pool was also reverse transcribed and Illumina sequencing library prep followed by 8-10 cycles of PCR using High Fidelity Phusion (NEB). As Msi1 concentration was increased, decreasing input RT reaction was required in the PCR. For instance, the highest Msi1 condition required 30-fold less input RT product than the no Msi1 condition. All libraries were barcoded in the PCR step, pooled together and sequenced one HiSeq 2000 lane.

#### SELEX-derived binding site used for validation

~~~
GGCUUCUUAAGCGUUAGUUAUUUAGUUCGUUUGUU
~~~

#### RBNS RT primer

~~~
GCCTTGGCACCCGAGAATTCCA
~~~

#### RNA PCR (RP1)

~~~
AATGATACGGCGACCACCGAGATCTACACGTTCAGAGTTCTACAGTCCGACGATC
~~~

#### Barcoded Primers

~~~
CAAGCAGAAGACGGCATACGAGAT–BARCODE–GTGACTGGAGTTCCTTGGCACCCGAGAATTCCA
~~~

### *In vivo* overexpression and whole mount mammary gland staining

Mice were given Dox (Sigma) via drinking water at 2 g/L. Mice were induced with Dox for 7 weeks unless otherwise indicated. Inguinal mammary glands were spread on glass slides, fixed in Carnoy’s fixative (6:3:1, 100% ethanol: chloroform: glacial acetic acid) for 2 to 4 hrs at room temperature, washed in 70% ethanol for 15 min, rinsed through graded alcohol followed by distilled water for 5 min, then stained in carmine alum overnight, washed in 70%, 95%, 100% ethanol for 15 min each, cleared in xylene and mounted with Permount.

### Quantitative PCR Analysis

Mouse mammary epithelial cells were prepared according to the manufacturer’s protocol (StemCell Technologies, Vancouver, Canada). Briefly, following removal of the lymph node, mammary glands dissected from 10-week-old virgin female mice were digested in EpiCult-B with 5% fetal bovine serum (FBS), 300 U/mL collagenase and 100 U/mL hyaluronidase for 8 hrs at 37°C. After vortexing and lysis of the red blood cells in NH_4_Cl, mammary epithelial cells were obtained by sequential dissociation of the fragments by gentle pipetting for 1-2 min in 0.25% trypsin, and 2 min in 5 mg/mL dispase plus 0.1 mg/mL DNase I (DNase; Sigma). Total RNA was isolated from mammary epithelial cells. Complementary DNA was prepared using the MMLV cDNA synthesis kit (Promega). Quantitative RT-PCR was performed using the SYBR-green detection system (Roche). Primers were as follows:

*P57* forward primer: 

~~~
GTTCTCCTGCGCAGTTCTCT
~~~

; *P57* reverse primer: 

~~~
GAGCTGAAGGACCAGCCTC
~~~

.

*Smad2* forward primer: 

~~~
TTTGCTGTACTCAGTCCCCA
~~~

;Smad2 reverse primer: 

~~~
TGAGCTTGAGAAAGCCATCA
~~~

*Smad3* forward primer: 

~~~
ACAGGCGGCAGTAGATAACG
~~~

;Smad3 reverse primer: 

~~~
AACGTGAACACCAAGTGCAT
~~~

*Smad5* forward primer: 

~~~
CTCATAGGCGACAGGCTGA
~~~

; *Smad5* reverse primer: 

~~~
AGATGGCCCCAGATAATTCC
~~~

*Tgfb1* forward primer: 

~~~
CAACCCAGGTCCTTCCTAAA
~~~

; *Tgfb1* reverse primer: 

~~~
GGAGAGCCCTGGATACCAAC
~~~

*Tgfb2* forward primer: 

~~~
TTGTTGAGACATCAAAGCGG
~~~

; *Tgfb1* reverse primer: 

~~~
ATAAAATCGACATGCCGTCC
~~~

*P21* forward primer: 

~~~
ATCACCAGGATTGGACATGG
~~~

; *P21* reverse primer: 

~~~
CGGTGTCAGAGTCTAGGGGA
~~~

*Hey1* forward primer: 

~~~
TGAGCTGAGAAGGCTGGTAC
~~~

; *Hey2* reverse primer: 

~~~
ACCCCAAACTCCGATAGTCC
~~~

*Msi2* forward primer: 

~~~
ACGACTCCCAGCACGACC
~~~

; *Msi2* reverse primer: 

~~~
GCCAGCTCAGTCCACCGATA
~~~

*K8* forward primer: 

~~~
ATCAAGAAGGATGTGGACGAA
~~~

; *K8* Reverse primer: 

~~~
TTGGCAATGTCCTCGTACTG
~~~

.

*K14* forward primer: 

~~~
CAGCCCCTACTTCAAGACCA
~~~

; *K14* Reverse primer: 

~~~
AATCTGCAGGAGGACATTGG
~~~

K18 forward primer: 

~~~
TGCCGCCGATGACTTTAGA
~~~

; K18 Reverse primer: 

~~~
TTGCTGAGGTCCTGAGATTTG
~~~

.

### Immunofluorescence on mammary gland sections

Mammary glands were fixed in 4% PFA, paraffin-embedded and 5-μm sections were used for immunofluorescence assay. Paraffin sections were microwave pretreated, and incubated with primary antibodies, then incubated with secondary antibodies (Invitrogen) and counterstained with DAPI in mounting media. The following antibodies were used: anti-K14 (Abcam), anti-K8 (Abcam), anti-E-cadherin (CST), anti-Msi2 (Novus Biologicals), anti-Hes1 (Abcam), anti-Slug (CST).

## Acknowledgements

We thank V. Butty, P. Reddien, P. Sharp, F. Soldner, J. Muffat, R. Weinberg, L. Surface, N. Spies, R. Friedman, M. Kharas for helpful discussions, R. Flannery for assistance with mouse colony maintenance, and D. Fu for assistance processing histology sections.

## Supplementary Figure Legends

**Supp. Figure 1. (A)** Distributions of the percent of tumors with non-silent mutations across cancer types in TCGA DNA sequencing data. Red and green triangles indicate values for Msi1 and Msi2, respectively. **(B)** Unsupervised hierarchical clustering of breast cancer tumors and matched controls, with overlaid sample labels, clinical markers and PAM50 subtypes.

**Supp. Figure 2. (A)** Unsupervised hierarchical clustering of gene expression from RNA-seq of breast cancer cell lines. (**B**) Fold-change in tumor-control pairs of TCGA breast cancer tumors for Msi1 and Msi2 across tumor subtypes. Msi1 shows a variable bimodal distribution of fold changes, while Msi2 is enriched in Luminal B tumors relative to Basal tumors. (**C**) Ratio of luminal to basal cancer cell line fold changes for Msi1, Msi2, Jag1 and Fn1.

**Supp. Figure 3. (A)** Quality control metrics for overexpression Ribo-Seq libraries. Left panel: percentage of reads mapped to genome, and the percentages of reads that are unique (“percent_unique”) and mapping to rRNA (“percent_ribo”) out of those mapped. Right panel: Percentage of reads mapping to exons (“percent_exons”), and out of those the percentage of reads in CDS regions (“percent_cds”), 3’ UTRs (“percent_3p_utr”), 5’ UTRs (“percent_5p_utr”). Percentage of reads mapping to introns (“percent_introns”) also shown. **(B**) Quality control metrics for knockout Ribo-Seq libraries, same format as (A).

**Supp. Figure 4. (A)** Top: Gel-shift MSI1 binding assay for Jag1 3’ UTR sequence 1. Kd estimate shown (15 nM) is average of 2 gel shifts. Bottom: Gel-shift for Jag1 3’ UTR sequence 1 mutant, where UAG sites mutated to UCC. Kd cannot be estimated (no binding to mutant could be detected.) **(B)** Top: Gel-shift MSI1 binding assay for Jag1 3’ UTR sequence 2. Kd estimate shown (9 nM) is average of 2 gel shifts. Bottom: Gel-shift for Jag1 3’ UTR sequence 2 mutant, where UAG sites are also mutated to UCC. Kd for mutant sequence was 649 nM.

**Supp. Figure 5.** Fold-change in Jag1 expression in Msi1 overexpression and double knockout samples for Ribo-Seq and RNA-Seq experiments.

**Supp. Figure 6. (A)** Immunofluorescence staining in mouse neural stem cells for MSI1 (red) and hnRNP A2/B1 (green). MSI1 shows predominantly cytoplasmic localization, while hnRNP A2/B1, a splicing factor, is predominantly nuclear. Confocal maximum Z intensity projections shown, 10 μm scale. (B) Western blot analysis for MSI1/2 and alpha-Tubulin (TUB) in total protein lysate (T), cytoplasmic protein lysate (C) and nuclear protein lysate (N) in control and Msi2 overexpressing cells.

**Supp. Figure 7. (A)** Distribution of MISO ΔΨ values in matched tumor-control pairs for Erbin (Erbb2ip) exon in light blue and Myo18a in dark blue. Right and left shifts from center (marked by dotted grey line at ΔΨ = 0) indicate tumor-enhanced and tumor-repressed splicing patterns, respectively. **(B)** Comparison of RNA fold changes in matched tumor-control pairs for Msi1 and Msi2 in Basal (left) and Luminal (right) tumors with ΔΨ values for Erbin and Myo18a exons. Points/triangles indicate luminal/basal tumor types determined by PAM50.

**Supp. Figure 8. (A)** Western blot for BT474 cells with control (shLuc) or Msi1/2 targeting hairpins. **(B)** Morphology of MCF7-Ras cells upon Musashi knockdown.

**Supp. Figure 9. (A)** Msi2 expression in mammary glands co-stained with basal cell marker K14 in control and Msi2 overexpressing mice. **(B)** Co-staining for luminal cell marker K8 and basal cell marker K14 in control (left) and Msi2 overexpressing (right) mice. **(C)** Staining for EMT marker Slug in control and Msi2 overexpressing mice. Scale bar: 50 μm.

